# Biological Reasoning with Reinforcement Learning through Natural Language Enables Generalizable Zero-Shot Cell Type Annotations

**DOI:** 10.1101/2025.06.17.659642

**Authors:** Xi Wang, Runzi Tan, Bo Wang, Simona Cristea

**Affiliations:** Department of Data Science, Dana-Farber Cancer Institute; Department of Computer Science, University of Toronto, Canada; Vector Institute for Artificial Intelligence, Toronto, Canada; Peter Munk Cardiac Center, University Health Network, Toronto, Canada; Department of Laboratory Medicine and Pathobiology, University of Toronto, Canada; Department of Biostatistics, Harvard T.H. Chan School of Public Health

## Abstract

Single-cell RNA-sequencing (scRNAseq) has reshaped biomedical research, enabling the high-resolution characterization of cellular populations. Yet cell type annotation, a process typically performed by domain experts interpreting gene expression patterns by manual curation or with specialized algorithms, remains labor-intensive and limited by prior knowledge. In addition, while reasoning large language models (LLMs) have demonstrated remarkable performance on mathematics, coding and general-reasoning benchmarks, their potential in scRNAseq analyses remains underexplored. Here, we investigate the advantages and limitations of employing DeepSeek-R1-0528, a recently developed open-source 671B-parameter reasoning LLM, for zero-shot scRNAseq cell type annotation. We find that DeepSeek-R1 prompted with a ranked list of 10 differentially expressed marker genes per cluster of single cells outperforms both its reasoning-enhanced, non-reasoning equivalent (DeepSeek-V3-0324) and GPT-4o in cluster-level annotations. At the level of single cells, DeepSeek-R1 prompted with the top 500 expressed genes in a cell outperforms its non-reasoning counterpart DeepSeek-V3, illustrating test-time scaling for bioinformatics tasks through natural language. Running DeepSeek-R1 in zero-shot classifier mode, with a prompt that presents a broad catalogue of cell type labels to choose from, improves its performance and generalizability across different datasets. On data curated by the expert model scTab (termed *in-domain* data), the DeepSeek-R1 classifiers perform better than the expert model scGPT and on par with the specialized cell genomics LLM C2S-Scale-1B, but lag behind scTab. On *out-of-distribution* data unseen by the two expert models, DeepSeek-R1 and its classifier versions generalize better and outperform the other models in the majority of the evaluated datasets. Notably, DeepSeek-R1 supports its cell type calls with interpretable textual biological rationales underlying its reasoning, providing a learning opportunity for researchers. Nevertheless, peak annotation performance remains modest, highlighting the intrinsic complexity of scRNAseq cell type annotation.

## 1. Introduction

Single-cell RNA-sequencing (scRNAseq) has transformed modern biology, enabling the study of gene expression in individual cells and revealing previously unrecognized cellular states. From immunology to developmental biology and precision oncology, scRNAseq approaches are now integral to addressing diverse research questions^1–6^. However, despite immense algorithmic progress within the past 10 years, with hundreds of advanced methods developed by the research community, cell type annotation remains a bottleneck step in scRNAseq bioinformatics pipelines. In practice, biomedical researchers often resort to manually annotating scRNAseq data by interpreting representative marker genes using their domain expertise. But, human experts can also be subject to bias in their annotations due to their specific expertise, experience level, or different perceptions of the granularity required for the task^7–9^.

Alternatively, *state-of-the-art* (SOTA) supervised methods or foundation models can accelerate cell type annotation. Yet, an inherent limitation of such approaches is that models can only recognize the cell identities present during training. As a result, novel or *out-of-domain (OOD)* cells are either forced into the closest known class or reported as “unknown”. Therefore, the true extent to which such algorithms generalize to novel datasets and offer practical utility for researchers as an end-to-end solution for cell type annotation remains unclear^10–15^.

LLMs have captured extensive attention for their capacity to reason through complex tasks using chain-of-thought (CoT), performing particularly well in mathematics, coding, or clinical decision-making^16–25^. DeepSeek-R1-0528 was recently introduced as an open-source general-purpose 671B-parameter reasoning model, specifically trained to strengthen its reasoning capabilities^16^. Reasoning models like DeepSeek-R1-0528 (referred to as DeepSeek-R1 or simply R1 from here onwards) can parse new problems at inference time, a concept known as test-time scaling, with little or no additional training^18^. This approach mirrors how human experts retrieve and synthesize their domain knowledge, suggesting that reasoning LLMs might contribute particularly well to tasks that rely on the dynamic interpretation and aggregation of complex data, such as scRNAseq cell type annotation.

As scRNAseq data presents itself in a quantitative format of gene-expression profiles, rather than a textual format, the potential of applying out-of-the-box existing generic LLMs to scRNAseq analysis tasks such as cell type annotation remains underexplored^7,26–31^. Attempts at using LLMs for cell type annotation include zero-shot cluster-level labeling with GPT-4 by *Hou et al. (2024)*,^7^ generating embedding representations via natural language prompts with CELLama^30^ or GenePT,^29^ as well as fine-tuning LLMs with scRNAseq data with Cell2Sentence and C2S-Scale^31,32^. Specifically, the *Hou et al. (2024)*^7^ paper showed how an early version of GPT-4 prompted with the top 10 differentially expressed marker genes per cluster can achieve near-expert accuracy and outperform existing SOTA expert algorithms such as singleR^33^. However, none of the existing works investigate the CoT reasoning capabilities of very recent generic reasoning LLMs such as DeepSeek-R1 for zero-shot scRNAseq cell type annotation, at both the cluster and single-cell levels.

We hypothesized that DeepSeek-R1-0528 is able to deploy its test-time logical reasoning abilities to interpret its pre-training biological knowledge and ultimately reliably annotate scRNAseq data, in a process conceptually similar to manual annotation by domain experts. Here, we tested this hypothesis by zero-shot prompting DeepSeek-R1 with representative marker genes for clusters or single cells, as either a stand-alone model or as a classifier contextualized with cell type labels to choose from. We focused specifically on assessing zero-shot cell type annotation capabilities, *i.e.* without relying on supervised fine-tuning, as supervised fine-tuning would require access to both bioinformatics expertise and labeled data, introducing practical real-world bottlenecks.

We compared DeepSeek-R1’s performance against non-reasoning LLMs, the expert models scTab^14^ and scGPT^15^, as well as the specialized cell genomics pretrained LLM C2S-Scale-1B^32^. We documented the advantages and limitations of annotating scRNAseq data with a general reasoning LLM across various tissues and datasets, and identified generalization challenges faced by the specialized models on unseen data^34^. Due to its reasoning nature, DeepSeek-R1 justified its annotations with biologically interpretable CoT, preserving the interpretability of manual marker-based annotation workflows and providing researchers with the opportunity to understand which genes were relevant for annotation and how they linked to the predicted label. With the amount of scRNAseq data increasing exponentially, reasoning LLMs such as DeepSeek-R1 can be leveraged to re-frame the cell type annotation problem altogether and identify the middle ground between generalizability and contextual expertise needed for annotating cells from novel experimental and biological setups.

## 2. Results

### 2.1. Reasoning with large language models for interpretable cell type annotations

Broadly, scRNAseq cell type annotation analyses adopt one of two main strategies: 1) a cluster-level approach, in which a cell type label is given to an entire cluster of cells by manual or reference-driven annotations, and 2) a single-cell level approach, in which each cell’s expression profile is mapped directly to a label, often using large classification models, reference-based tools or, more recently, foundation models. Despite the open-source availability of increasingly sophisticated cell type annotation algorithms, researchers still rely on manual inspection of canonical marker genes, regarded in practice by most molecular biologists and medical professionals as the “gold standard” to ensure biological interpretability. However, manual annotation remains a labor- and time-intensive process.

Here, we propose an alternative cell type annotation approach that leverages the general-purpose reasoning LLM DeepSeek-R1^16^ (Fig. 1A). We adopted a prompt engineering technique similar to that of *Hou et al. (2024)* and *Lu et al. (2024)* that exploits the LLM’s zero-shot cell type annotation capabilities at inference^7,31^. We prompted the LLM with either a cluster’s top 10 differentially expressed marker genes, or a single cell’s top 500 expressed genes, along with relevant metadata (*e.g.*, species or tissue), and asked it to identify the granular cell type label. DeepSeek-R1 responded with CoT reasoning, culminating in an annotated cell type label (Fig. 1B, C). Furthermore, we investigated the impact of prompt engineering on cell type annotation by comparing “long” prompts against “short” prompts asking for more concise responses.

**Figure 1.**
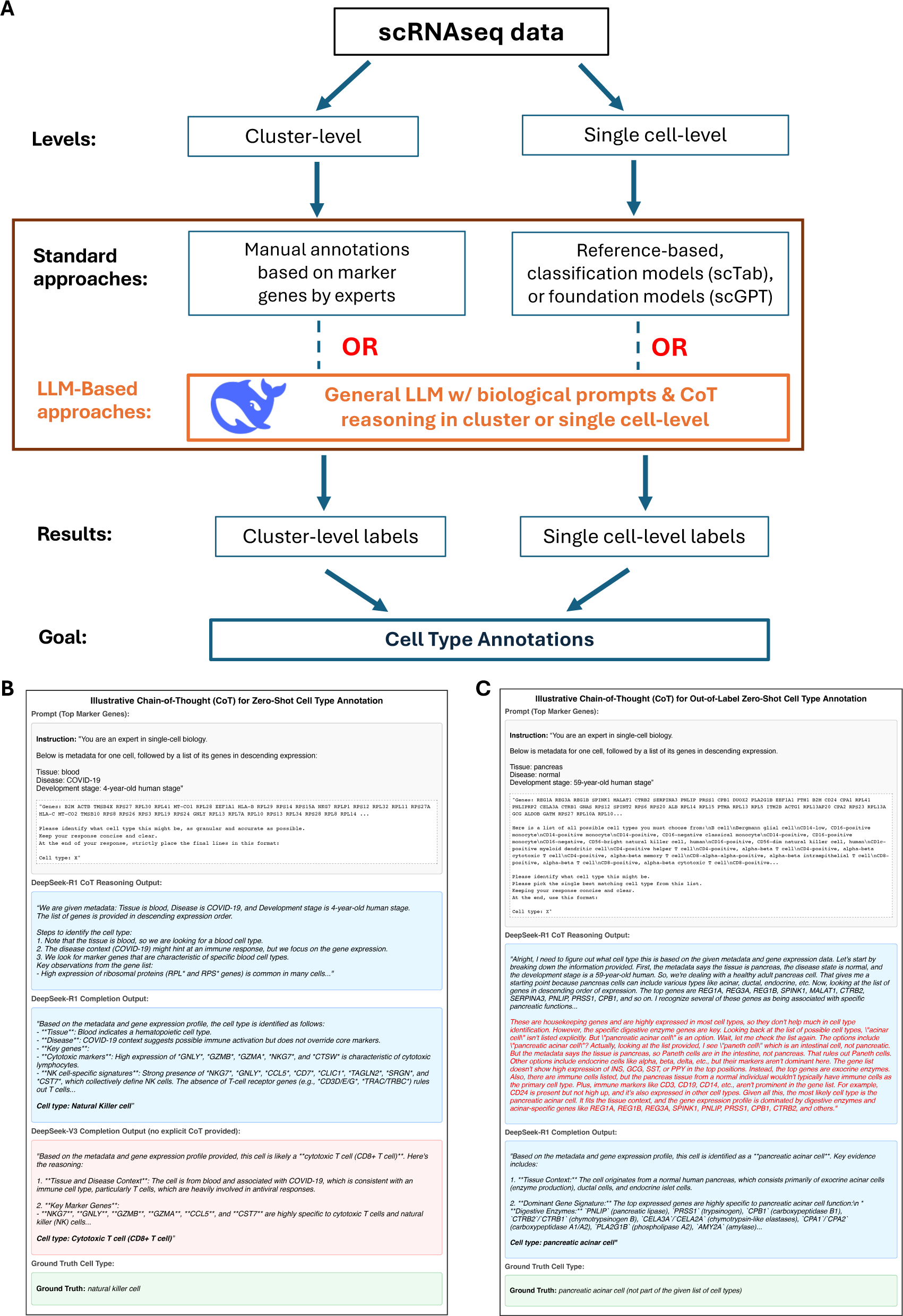
Overview of the proposed reasoning LLM-based cell type annotation framework and illustrative chain-of-thought biological LLM reasoning. (A) Two parallel strategies for scRNAseq cell type annotation: cluster-level and single-cell level. Standard pipelines rely on a combination of manual marker gene inspection, reference-based methods or deep learning classifiers. In contrast, this approach employs a general-purpose reasoning LLM (DeepSeek-R1) to process ranked gene signatures in either a cluster-level (top 10 differentially expressed genes) or single-cell level context (top 500 highest expressed genes). (B) Example chain-of-thought outputs from DeepSeek-R1 and DeepSeek-V3, where rank-ordered genes and basic metadata guide zero-shot labeling of a single cell. DeepSeek-R1’s reasoning links key genes to known biological functions, ultimately predicting “Natural Killer cell,” matching the ground truth label of a natural killer cell, while the non-reasoning model DeepSeek-V3 predicted the wrong cell type. (C) Additional example chain-of-thought output for single-cell level annotation, in which DeepSeek-R1 is prompted to choose a cell type label from a list of pre-specified unique cell types employed by the model scTab^14^ within its classification, essentially turning LLM cell type annotation into a multi-class classification problem. However, in this case, the ground truth cell type is not among the list of unique labels provided. Via its reasoning, DeepSeek-R1 correctly recognizes the limitation of the provided list of cell types and proposes the correct cell type outside of the original list.

### 2.2 LLM reasoning enhances cluster-level cell type annotations

Our cluster-level analysis used the 1,130 single-cell clusters from the work by *Hou et al. (2024)*^7^ for benchmarking. This aggregated dataset originates from different studies encompassing multiple tissues and cell types from human and mouse, including lung, skin, blood, prostate, fetal development, and many others (Fig. 2A). For performance evaluation and benchmarking, we adapted scTab’s evaluation framework^14^ with an additional label-matching step, as done in *Hou et al. (2024)*^7^. Specifically, for each cluster, given a label predicted by a model and a ground truth label provided by the original dataset, we first matched both labels to their corresponding Cell Ontology (CL) database terms^35^, to remove potential ambiguities in LLM outputs. Then, using Ubergraph^36^, we compared the ground truth label first with the LLM output matched against CL, then further with all its child descendants in the ontology tree. If a match was recorded between the ground truth label and either the LLM label or the label of any of its descendants, the prediction was recorded as TRUE. Otherwise, the match was unsuccessful, and the prediction was recorded as FALSE. Lastly, we aggregated these TRUE/FALSE predictions across all the 1,130 tested clusters (Methods).

**Figure 2.**
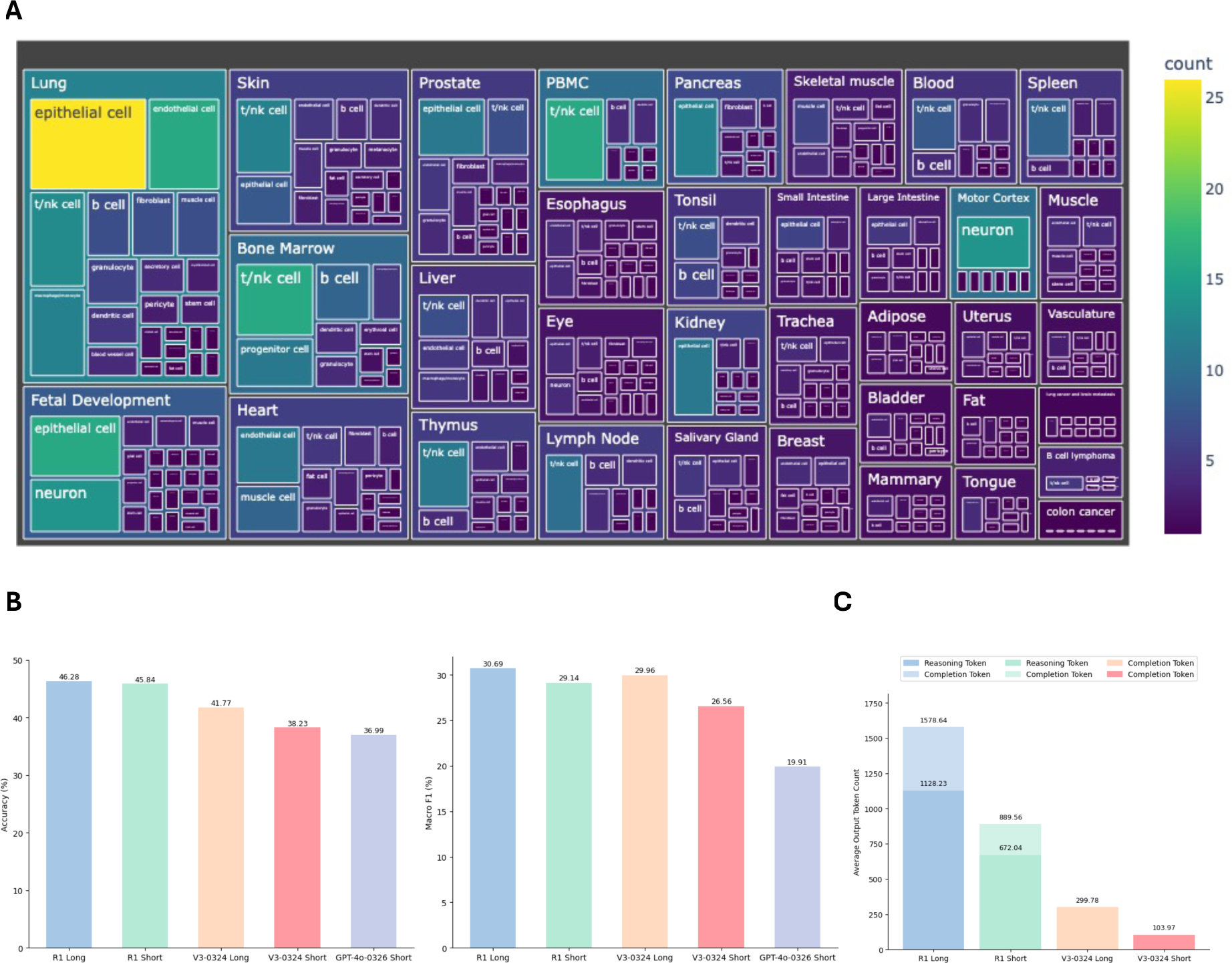
Benchmark performance of DeepSeek-R1 on cluster level cell type annotation against DeepSeek V3 and GPT-4o. (A) Treemap plot showing the cluster-level dataset composition across cell types and tissues; the color scale and the area of each box indicate the number of clusters per each unique tissue and cell type combination. (B) Bar plots comparing the accuracy and Macro-F1 score of the tested models on cluster-level cell type annotation across the 1,130 clusters. GPT-4o-0326 was tested using the same short prompts as R1 and V3. (C) Bar plots showing the average token count for cluster-level cell type annotation, averaged across clusters. For reasoning models, the darker color represents the reasoning token count (unique to reasoning models, representing the computational steps during the internal thought process), and the lighter bar shows the completion token count (representing all the final generated answer text); for non-reasoning models, the bar shows the completion token count.

As the *Hou et al. (2024)* study showed that a now-outdated version of GPT-4 was superior to SOTA expert models such as SingleR^33^, scType^37^, and CellMarker 2.0^38^ for cluster-level cell type annotation, in our cluster-level analysis we benchmarked DeepSeek-R1 only against the contemporary GPT-4o (version 2025-0326) and DeepSeek-V3-0324. DeepSeek-V3-0324 is an updated version of DeepSeek-V3, the instruction-tuned and Reinforcement Learning with Human Feedback (RLHF)-fine-tuned version of the DeepSeek-V3-base model (unavailable at the time of testing). Notably, DeepSeek-V3-0324 incorporates reasoning enhancements from the first version of DeepSeek-R1 (DeepSeek-R1-0120) via improved post-training and RL insights. However, while both DeepSeek-R1 and DeepSeek-V3-0324 can explain their answers step-by-step, the R1 models were built from the ground up as reasoning models, while V3-0324 learned to reason by distilling DeepSeek-R1-0120’s CoT^16^. Therefore, the R1 models are placed in the Arena’s reasoning tier, while V3-0324 is considered reasoning-enhanced but classified as non-reasoning. In contrast, GPT-4o has not been explicitly trained with techniques like long-form CoT distillation or targeted RLHF that reward internal reasoning steps and is considered a full non-reasoning model.

Our evaluations revealed that both DeepSeek models outperformed GPT-4o in annotation accuracy, with the highest cluster-level accuracies of 46.28% and 45.84% obtained by DeepSeek-R1 with the long reasoning and short prompt respectively, and the lowest accuracy of 36.99% obtained by GPT-4o (Fig. 2B). DeepSeek-R1’s accuracy also consistently surpassed DeepSeek-V3-0324 under the same prompt (46.28% vs 41.77% for long prompts and 45.84% vs 38.23% for short ones), with Macro-F1 evaluations following a similar trend (Fig. 2B; Supplementary Table S1).

We further examined how different prompting strategies affected test-time computation by investigating the average token count per query (Fig. 2C). For DeepSeek-R1, the output includes internal *reasoning tokens* (representing the model’s step-by-step thought process before generating the final answer) along with the final output tokens intended for the user. The *completion token* count for R1 is the sum of both reasoning and final output tokens. In contrast, V3-0324 does not have explicit internal reasoning tokens; rather its output consists solely of *completion tokens*. We found that, on average, R1 used far more tokens than V3-0324: 5.27 times more for the long prompt, and 8.56 times more for the short prompt. The absolute 0.44% increase in R1’s accuracy when using a long versus a short prompt came at the cost of average reasoning and completion token increases of 1.68 and 1.77-fold respectively, while for V3-0324, an absolute 3.54% accuracy gain came at the cost of 2.88-fold more completion tokens.

Altogether, these results show that: 1) enhanced LLM logical reasoning during inference, particularly longer test-time, translates into improved biological interpretation of cluster-level differential gene signatures compared to lack of reasoning, and 2) the performance benefit of full reasoning (R1) comes at a high token cost compared to learned reasoning capabilities (V3-0324), and similarly for long versus short prompts, in line with recent reports of sub-optimal token consumption of full reasoning models^39–43^. Lastly, the performance metrics reported here were lower than those of *Hou et al. (2024)* for similar models ran on the same cluster-level data because of the different scoring strategy employed by *Hou et al (2024)*, in which they also considered partial matches between ground-truth and predicted labels.

### 2.3. DeepSeek-R1 and its variants overperform expert models on out-of-domain data and scTab excels on known data

Compared to cluster-level annotation, the annotation of individual cells is more challenging due to higher granularity and lower signal-to-noise ratio. Here, using the same evaluation methodology as for the cluster-level analysis, we benchmarked DeepSeek-R1 first against the non-reasoning model DeepSeek-V3, and further against three specialized models: the two expert models scTab^14^, a multi-class classifier, and scGPT^15^, a scRNAseq foundation model, and the single cell genomics pretrained LLM C2S-Scale-1B^32^. For benchmarking, we chose DeepSeek-V3 as it is the backbone for both DeepSeek-V3-0324 and DeepSeek-R1 and it does not incorporate any reasoning objective, scTab and scGPT as recent SOTA models for scRNAseq single cell annotation, and C2S-Scale-1B as a representative tool for LLM scRNAseq cell type annotation using similar gene lists prompts. C2S-Scale-1B is based on the Pythia-1B architecture and fine-tuned using the Cell2Sentence^31^ framework on a wide array of scRNAseq datasets.

Our benchmarking utilized four datasets derived from CellXGene: 1) An *in-domain validation dataset*: 10,000 cells randomly subsampled from the 3.5 million cells used as curated validation data by scTab^14^; 2) Similar to 1), an *in-domain test dataset* consisting of 10,000 cells randomly subsampled from scTab’s curated test dataset of 3.4 million cells; 3) An *out-of-domain (OOD) random dataset*: 10,000 cells randomly subsampled from the 7.8 million cells added to CellXGene after May 15^th^ 2023, representing data not seen by either scTab or scGPT during training or testing^44^ (Fig. 3A); 4) Similar to 3), an *OOD balanced tissue dataset*, with a different subset of randomly sampled 10,000 cells from the same set of 7.8 million cells, this time covering eight different tissues, with 1,250 sampled cells per tissue (Methods). As the fine-tuning cut-off date for C2S-Scale-1B was later than the release dates of the *OOD* datasets, the *OOD* cells were likely explicitly seen by the model prior to inference, especially given C2S-Scale-1B’s fine-tuning strategy with cell sentences consisting of lists of marker genes and corresponding ground-truth labels sourced from CellXGene^44^ and the Human Cell Atlas^45^. In contrast, even though DeepSeek-R1 could have in principle also encountered material related to the *OOD* datasets during training, the DeepSeek pre-training corpus is almost exclusively open-access text and excludes scRNAseq matrices or preprocessed marker-gene tables. Therefore, any potential exposure would have likely been limited to statements linking selected canonical marker genes to cell types, rather than to the verbatim comprehensive gene lists used in our prompts.

**Figure 3.**
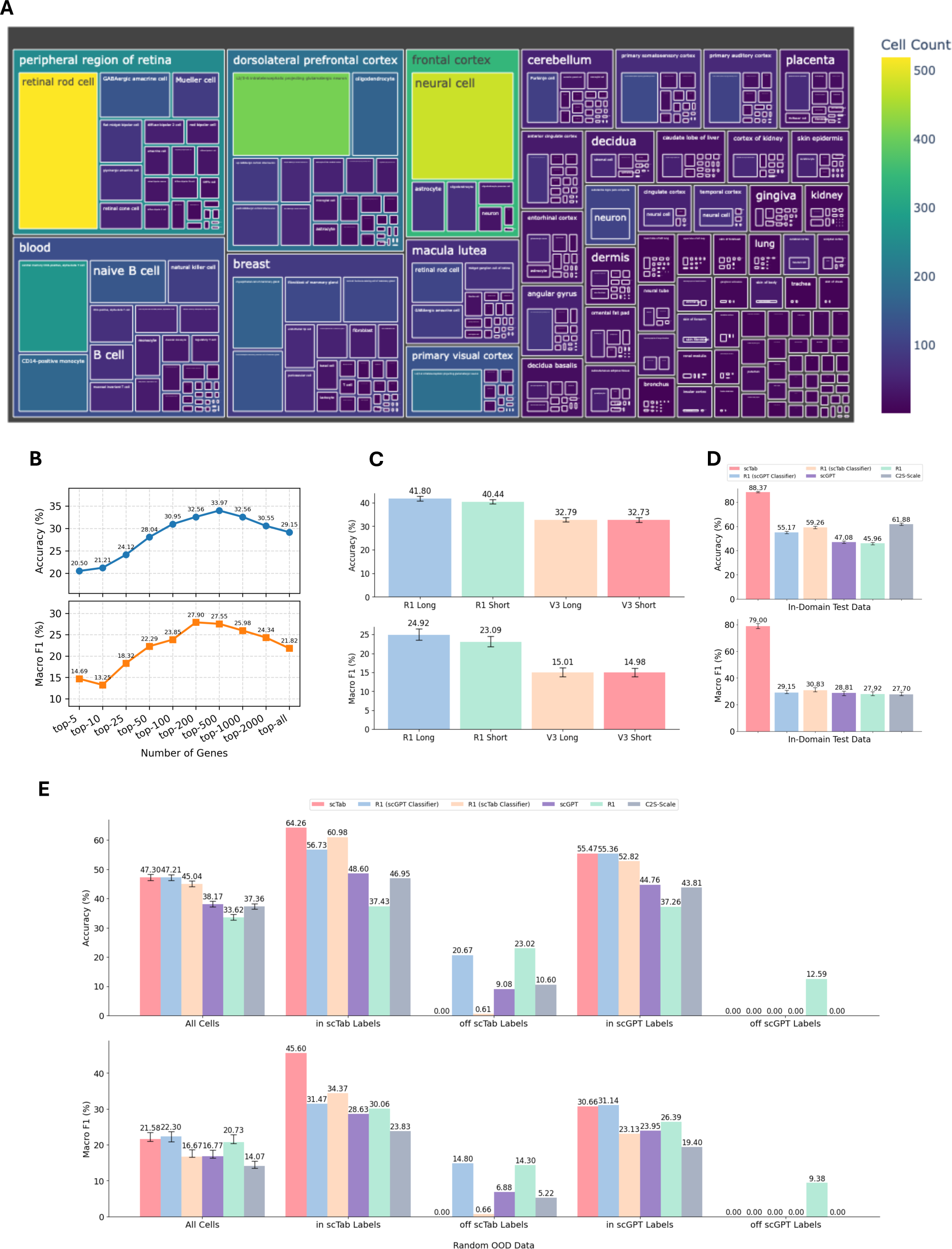
Benchmark performance of DeepSeek-R1 on single-cell level cell type annotation against DeepSeek-V3 and specialized cell type annotation models on *in-domain* and *OOD* random datasets. (A) Treemap plot showing the tissue and cell type composition of the 10,000 cells in the *OOD random dataset*, added to the CellXGene database after May 15^th^, 2023, the cutoff for both scGPT and scTab development. The color scale and the area of each box indicate the number of cells per each unique tissue and cell type combination. (B) DeepSeek-R1’s accuracy and Macro-F1 score as a function of *N*, the number of top expressed genes included in the prompt, based on a subset of 1,000 randomly sampled cells from the *OOD* random dataset. (C) Accuracy and Macro-F1 score of DeepSeek-R1 and DeepSeek-V3 on the *in-domain* validation dataset consisting of 10,000 randomly sampled cells. Long and short denote different prompting strategies. Due to DeepSeek-V3 model availability in the API, this benchmarking was run with the top 100 highly expressed genes in the prompt. (D) Accuracy and Macro-F1 score of DeepSeek-R1, the two classifier versions of DeepSeek-R1 using the cell type label set of either scTab or scGPT, the two expert models scTab and scGPT and C2S-Scale-1B on the *in-domain* test dataset consisting of 10,000 randomly sampled cells. (E) Performance of the same six models as in (D) on the *OOD* random dataset on all cells, as well as split by whether the ground-truth cell type labels were within scTab’s or scGPT’s labels.

Before benchmarking, we first identified the optimal number N of highly expressed genes per cell to include in DeepSeek-R1’s prompt, using 1,000 randomly subsampled cells from the *OOD* random dataset for multiple values of N, ranging from 5 (corresponding to the top 5 most highly expressed genes per cell) to the full set of all genes expressed in a cell (Fig. 3B). We found that providing the ranked 500 highest expressed genes in the prompt yielded the highest accuracy and the second-highest Macro-F1 score, and we therefore set N = 500 as prompt input for subsequent analyses. For the *in-domain* and random *OOD* datasets, we computed confidence intervals (CI) for performance metrics, as well as pairwise comparisons of model performance (Methods).

#### 2.3.1. In-domain benchmarking

We first evaluated the performance of the reasoning model DeepSeek-R1 against its non-reasoning counterpart DeepSeek-V3 on the *in-domain validation dataset* using both long and short prompts (Methods). R1 outperformed V3 in both accuracy and Macro-F1 score, while long prompts performed similarly to short prompts, with a significant boost in long prompting for R1 (Fig. 3C; Supplementary Fig. 1). We then benchmarked DeepSeek-R1 with long prompts against scTab, scGPT and C2S-Scale-1B. Recognizing the classifier nature of the expert models scTab and scGPT, we also evaluated R1 in classifier mode by constraining the model to choose a single label from a pre-defined set of labels given in the prompt (Methods). scTab imposed stricter data curation restrictions than scGPT, such as requiring any cell type to have at least 5,000 unique cells present in at least 30 donors. In consequence, scTab’s label set consisted of 164 cell types, as compared to 593 for scGPT.

For *in-domain* testing, we used scTab’s test data. As expected, scTab outperformed all models on this curated dataset, followed by the C2S-Scale and DeepSeek-R1 classifiers with scTab and scGPT labels, with the classifiers performing better than C2S-Scale on Macro-F1 scores (Fig. 3D). scGPT and the unconstrained DeepSeek-R1 scored similar accuracy (Supplementary Fig. 1), highlighting the capacity of general reasoning LLMs to annotate single cells at similar levels to SOTA expert models, even in controlled settings where all ground truth cell type labels are within the dictionaries of the expert models.

#### 2.3.2. Out-of-domain (OOD) random sampling benchmarking

When tested on 10,000 cells randomly downloaded from CellXGene after May 15^th^, 2023, and unseen by either of the two expert models, scTab and R1 with scGPT classifier scored statistically indistinguishable highest accuracy (Fig. 3E, upper panel labeled *All cells*; Supplementary Fig. 1). scTab’s random *OOD* accuracy dropped by 46% relative to *in-domain*, and C2S-Scale’s accuracy dropped by 40%. In contrast, R1’s performance was more robust *in-domain* and *OOD*, with relative drops of 14% for R1 with scGPT classifier, 24% for R1 with scTab classifier, and 27% for unconstrained R1. Similarly, scGPT’s accuracy only dropped 19% *OOD* relative to *in-domain*. The Macro-F1 score was highest for R1 with scGPT classifier labels, followed by scTab and unconstrained R1 (Fig. 3E, lower panel labeled *All cells*). The difference in ranking between accuracy and Macro-F1 score reflects different performances on more frequent and less frequent cell types, with models scoring higher on Macro-F1 generally having a more balanced performance across cell types of various frequencies.

The change in the models’ performance on *OOD* relative to *in-domain* data reflects variations in batch effects, data distribution and curation, annotation granularities and unseen cell types. Since the *OOD* cells were novel to scTab and scGPT, we investigated how much of the models’ decrease in performance was due to a lack of label overlap (*in-label* and *off-label* cells). As expected, scTab’s accuracy rebounded to 64.26% when only assessing on its labels and its Macro-F1 score to 45.60%, closely followed by R1 in classifier mode with scTab labels (60.98%), which nevertheless scored lower Macro-F1 (34.37%, Fig. 3E, panels labeled *in scTab Labels*). When evaluating only on cell types among scGPT’s labels, both scTab and R1 classifier with scGPT labels performed best (55.47% and 55.36% accuracy, Fig. 3E, upper panel labeled *in scGPT Labels*). For the subset of cells with labels not among scTab’s labels, R1 with scGPT classifier and unconstrained R1 had the highest accuracy and Macro-F1 scores, substantially outperforming scGPT, despite the expert model and the R1 classifier having access to the exact same set of labels (Fig. 3E, panels labeled *off scTab Labels*). The non-zero performance of R1 with scTab classifier was a consequence of the rare situations of R1 proposing a cell type label outside its instruction list from the prompt (3.79% of cases for R1 with scTab labels, and 0.63% for R1 with scGPT labels), and that prediction being correct (Fig. 1C). On cells off scGPT labels, unconstrained R1 was the only model with non-zero performance (Fig. 3E, panels labeled *off scGPT Labels*), underscoring DeepSeek-R1’s adaptability for labeling less commonly encountered cell types.

In summary, on the *OOD* random data, R1 in classifier mode with scGPT labels emerged as the most reliable overall cell type annotation strategy, consistently scoring high in most settings. The model also showed comparable accuracy and Macro-F1 scores when tested on the two sets of curated cells (164 cell type labels for scTab and 593 for scGPT) and performed on par with unconstrained R1 on cells off scTab labels. We note that the peak performance of all tested models was modest on the *OOD* random data, reflective of the realistic situation of employing LLMs or expert models for zero-shot annotating scRNAseq data. Even though all cells in this dataset had been QCed and assigned ground truth labels by their respective studies, this data has not undergone additional cell-type-targeted curation, and all cell types were considered before sampling, regardless of their relative frequencies and granularities, in contrast to scTab’s *in-domain* data.

#### 2.3.3. OOD balanced tissue benchmarking

To more accurately mimic real-world scenarios that researchers might encounter as daily bioinformatics tasks, we created an *OOD balanced tissue* dataset, in which we sampled 1,250 cells from eight selected tissues (blood, peripheral region of retina, kidney, breast, lung, trachea, pancreas and cerebellum), amounting to 10,000 cells (Fig. 4A; Supplementary Table S2; Methods). On this aggregated data, the R1 classifiers were the most accurate models across all meaningful settings (Fig. 4B, upper panels). R1 with scGPT labels performed best when tested across all cells, on cells off scTab labels and on cells within scGPT labels, while R1 with scTab labels scored highest when restricting the evaluation only to cells within scTab labels. The unconstrained R1 model emerged as top performing by Macro-F1 scoring in all scenarios except when restricting to scTab labels, further demonstrating how DeepSeek-R1 can generalize and reliably zero-shot annotate a wide variety of cell types of various frequencies and with widely different phenotypes.

**Figure 4.**
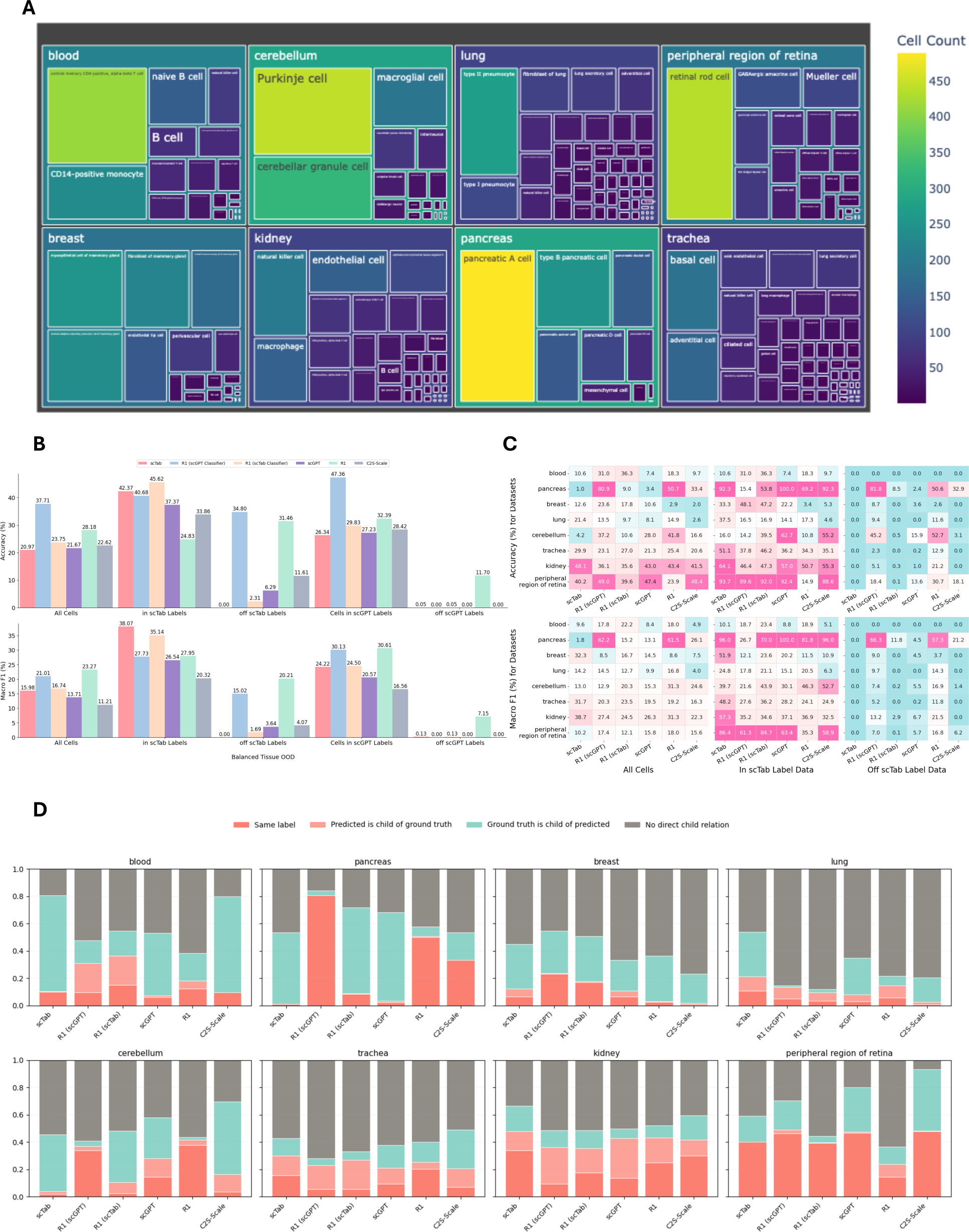
Benchmark performance of DeepSeek-R1 on single-cell level cell type annotation against specialized cell type annotation models on the *OOD* balanced tissue dataset. (A) Treemap plot showing the tissue and cell type composition of the 10,000 cells belonging to eight selected tissues (1,250 cells per tissue) in the *OOD* balanced tissue dataset used for benchmarking, added to the CellXGene database after May 15^th^, 2023, the cutoff for both scGPT and scTab development. The plot depicts the balanced dataset of 10,000 cells belonging to eight selected tissues, with each tissue contributing 1,250 cells. The color scale and the area of each box indicate the number of cells per each unique tissue and cell type combination. (B) Performance of the same six models as in Fig.3 D and E, with the same figure layout. (C) Accuracy and Macro-F1 score of the same six models as in (B), split by the eight selected tissues, as well as by whether the ground-truth cell type labels were part of scTab’s labels. (D) Stacked barplots showing the distribution of cell-level annotations across all six models and all eight tissues, split by whether the predicted and the ground truth label either matched perfectly or predicted was a child of ground truth (recorded as correct predictions), or ground truth was a child of predicted (incorrect prediction), or there was no direct child-parent relation between predicted and ground truth (also incorrect). For the lung tissue dataset, 382 of the 1,250 cells had no match of the ground truth label in Cell Ontology and were therefore excluded from accuracy (B and C) and frequency (D) evaluations.

To better understand why the *OOD* balanced-tissue benchmark was more challenging for the specialized models, especially scTab, we investigated the label overlap between the labels in these datasets and scTab’s label set (Supplementary Fig. S2). While scTab covered some lineages well, with 0.08% of blood cells, 24.96% of kidney cells, 41.44% of trachea cells, and 42.86% of lung cells falling outside its vocabulary, it missed a larger fraction of the remaining four tissues, with 57.04 % of retina, 62.24% of breast, 74.08% of cerebellum and virtually the entire pancreas dataset (98.48%) falling outside of scTab’s 164 labels. In total, 50.44% of cells with a ground truth Cell Ontology match (Supplementary Fig. S3) were off label for scTab, despite all eight tissues included in scTab’s classification. This discrepancy might have happened either because the newly added datasets included novel cell types that scTab’s datasets did not include, or because these cell types were not large or frequent enough to pass scTab’s strict curation criteria.

Further detailed breakdown of tissue-specific performance showed interesting patterns for the different models (Fig. 4C). For blood cells, the two DeepSeek-R1 classifiers yielded the best performance in both accuracy and Macro-F1 score, while scTab and scGPT performed much more poorly (Fig. 4C, *All Cells*), despite almost complete cell type label overlap (99.92%) with scTab and blood being the most frequent tissue in scTab’s training set. A closer inspection of the cell-level predictions (Fig. 4D; Supplementary Table S3) showed that the specialized models, especially scTab and C2S-Scale-1B, often missed the right granularity to match the ground-truth data (*e.g.* predicting “leukocyte” for a ground-truth label of “naïve B cell”). This suggests generalization challenges for the specialized models on novel datasets, despite having access to the right labels and having encountered a large number of similar cells during training. In contrast, when given specific indications to choose from either scGPT’s or scTab’s set of labels, DeepSeek-R1 in classifier mode correctly labeled a much larger fraction of cells, with even greater granularity than the ground truth data, providing additional biological information.

For pancreas, scTab’s training corpus only listed six non-specific and infrequent cell types: B cell, T cell, endothelial cell, mast cell, mature NK T cell, and plasma cell (Supplementary Table S4). In contrast, the pancreas *OOD* dataset analyzed in this study^46^ consisted of ten highly specific pancreatic cell types: pancreatic A cell, pancreatic D cell, pancreatic ductal cell, pancreatic PP cell, type B pancreatic cell, endothelial cell, mesenchymal cell, pancreatic acinar cell, pancreatic endocrine cell, and pancreatic epsilon cell. Endothelial cells, representing less than 5% of all pancreatic cells, were the only common cell type between the two datasets, while acinar cells represent 80-85% of the entire pancreatic tissue mass. Due to almost very low cell type label overlap (1.52%), scTab and the DeepSeek-R1 scTab classifier performed poorly on this dataset (Fig. 4C). In contrast, while scGPT’s performance was also poor (3.4%), contextualizing DeepSeek-R1 with scGPT’s labels increased its accuracy to 80.9% (compared to 50.7% for unconstrained R1), further suggesting generalization struggles of expert models. Closer inspection showed that the predictions of the two expert models scTab and scGPT were too general, borrowing similar cell types from other tissues (Supplementary Table S3, Fig. 4D).

On the breast tissue, DeepSeek-R1 in scGPT classifier mode showed the highest accuracy, while scTab had the highest Macro-F1 score (Fig. 4C). The performance of unconstrained R1 was particularly poor in accuracy (2.9%), due to granularity issues such as predicting “myoepithelial cell” instead of a ground truth of “basal myoepithelial cell” or mistaking the two lineages basal and luminal (Supplementary Table S3). The lung data showed highest accuracy for scTab and highest Macro-F1 score for unconstrained R1, likely a consequence of high cell type label overlap with scTab (Supplementary Fig. S2), as well as many lung cells used in scTab’s training. For cerebellum, the unconstrained R1 scored highest in both accuracy and Macro-F1 score, with the overwhelming majority of their correct predictions having the exact same label as the ground truth data, and very few incorrect predictions of lower granularity (Fig. 4D, Supplementary Table S3). For trachea and kidney, scTab yielded the highest accuracy and Macro-F1 score (Fig. 4C), likely also a consequence of high cell type label overlap (Fig. S2). Lastly, results on the peripheral region of retina tissue showed highest accuracy of R1 in scGPT classifier mode (Fig. 4C) and high percentages of perfect matches for all models except unconstrained R1 (Fig. 4D), even though unconstrained R1 was the model with the highest Macro-F1 score.

Taken together, these analyses on the *OOD balanced tissue* dataset revealed that balancing the tissue distribution and assessing cell type annotation performance separately for each tissue is essential for understanding model performance. Overall, the DeepSeek-R1 model demonstrated superior adaptability, especially in classifier mode when contextualized with appropriate label constraints from the large scGPT label vocabulary.

## 3. Conclusion and Discussion

Formulating the cell type annotation problem as a benchmarking task is inherently difficult. A necessary ingredient for successful cell typing is accurately labeling biological knowledge as either known or novel. In practice, some cell types are frequent, generic and display a distinctive marker profile, making them relatively straightforward to annotate regardless of the strategy employed (manually investigating a list of marker genes, querying an LLM, or running a specialized cell type annotation model). On the contrary, other cell types are infrequent across donors, their functionality is ambiguous, and their gene expression profiles are non-specific. Annotating such cells turns out to be difficult regardless of the approach used. Moreover, researchers often disagree on the most appropriate ground truth annotation label, or even whether such groups of cells indeed represent a novel cell type or are better characterized as an alternative cellular state of an already-existing cell type^47^. Nevertheless, rare and ambiguous cell types are crucial for novel biological discoveries^48^. Such aspects turn cell type annotation into a complex problem for which even evaluating the quality of a given solution, is challenging and subject of on-going debate^47^. Therefore, the ideal real-world cell type annotation procedure should not only be accurate, but also versatile and adaptable to different scenarios, some of which are hard to capture *a priori* as a set of pre-defined rules.

Building on this rationale, we hypothesized that recently developed general-purpose reasoning LLMs could have the potential to positively contribute to the scRNAseq cell type annotation toolbox, by striking an interesting balance between accuracy and discovery. To this end, we examined the feasibility of employing DeepSeek-R1-0528 to perform both cluster-level and single-cell level annotations in scRNAseq data in a zero-shot setting, without specialized fine-tuning. Running the LLMs zero-shot is key, as fine-tuning requires both existing labeled data and expert bioinformatics expertise, which can be a bottleneck in real-world situations.

By prompting DeepSeek-R1 with a list of ranked marker genes, we assessed its capacity to identify cell types through interpretable CoT reasoning that captures canonical markers, biological functions, and tissue-specific knowledge. We found that LLM reasoning enhanced cell type label prediction at both cluster and single-cell levels. In single-cell datasets, our results revealed that running DeepSeek-R1 in classifier mode, with its prompt contextualizing a large set of cell type labels to choose from, improved its performance, leading to overall superior adaptability and generalizability across tissues and datasets. When comparing DeepSeek-R1 and its classifier variants with the three specialized models scTab, scGPT and C2S-Scale-1B on the curated scTab *in-domain* data, general LLMs performed better than scGPT, while lagging behind scTab and C2S-Scale-1B. However, on random *OOD* data unseen by the expert models, the DeepSeek-R1 scGPT classifier performed on par with scTab, outperforming the other models. The expert models faced generalization challenges on unseen data^34^ and were outperformed by DeepSeek-R1 and its classifier versions on a separate *OOD* dataset consisting of cells from eight relatively common tissues. Notably, half of the cells in this *OOD* balanced tissue dataset fell outside scTab’s 164-label vocabulary, highlighting how strict curation criteria can inadvertently exclude biologically relevant but less frequent cell types, leading to decreased generalization capabilities.

Our study demonstrated that annotating scRNAseq data with general reasoning LLMs is reliable and interpretable. Employing such models for scRNAseq cell type annotation will allow the single-cell community to directly benefit from the AI foundation models and biological reasoning innovation wave^49^, with new SOTA general LLMs released very frequently. In addition, DeepSeek-R1’s interpretability and the biological context of its predictions can provide a transparent rationale to cell type annotation similar to manual marker-based annotations. In some situations, DeepSeek-R1 outputted correct predictions of higher granularity than ground-truth labels, reframing the cell type annotation process as a learning opportunity for researchers. Its adaptability counteracted the traditional limitation of tissue-specific training of expert models, allowing on-the-fly generalization to real-world situations and potentially revealing novel cellular biology.

At the same time, our study demonstrated that scRNAseq cell type annotation remains a challenging problem to evaluate and solve. Peak cluster-level and single-cell-level performance was overall modest for any dataset which was not heavily curated by removing infrequent cell types. This happened because real-world randomly-drawn unseen data remains difficult to annotate at the level of granularity proposed by the original study. Some proposed cell type labels are by design very specific, with potentially highly similar expression profiles to other cell types, making accurate differentiation challenging. Additionally, the evaluation criterion employed here was stringent: we required the predicted label to be at least as granular as the ground-truth label, leading to lower performance measures than reported elsewhere.

As recent AI agentic workflows that use LLMs for cell type annotation have been shown to perform better than the models alone^50,51^, we hypothesize that the entire scRNAseq cell type annotation process can be automated with a suite of specialized agents orchestrated by an LLM reasoning “brain” (Fig. 5). For example, the Data Ingestion Agent can take raw scRNAseq data and run basic quality checks. Next, the Clustering Agent can sort cells into subclusters, while an Annotation Agent can assign preliminary labels at either the cluster level or individual cell level. After that, a Quality Control (QC) Agent, backed by an Ontology Agent, a Label Verifier Agent, and a Reasoning Verifier Agent, can examine the given labels against known marker references and ontology databases. At any point, the orchestrator “brain” can call for deeper checks or bring in human expertise. Finally, an Output Agent can package the refined annotations, key performance metrics, and a summary of how the decisions were made for researchers to evaluate. Because DeepSeek-R1 is flexible and requires no domain-specific training, it could serve as the central orchestrator brain, iteratively integrating and verifying knowledge from the individual sub-agents with minimal human intervention. This multi-agent system is both interpretable and adaptable: the actions and rationale of each sub-agent remain clearly documented, preserving the transparency associated with manual marker gene review, and the orchestrator’s reasoning capabilities allow it to readily handle new *OOD* cell types. We anticipate that such automations could substantially accelerate the rapidly evolving landscape of single-cell research, boost efficiency, and allow scientists to focus on deeper biological questions.

**Figure 5.**
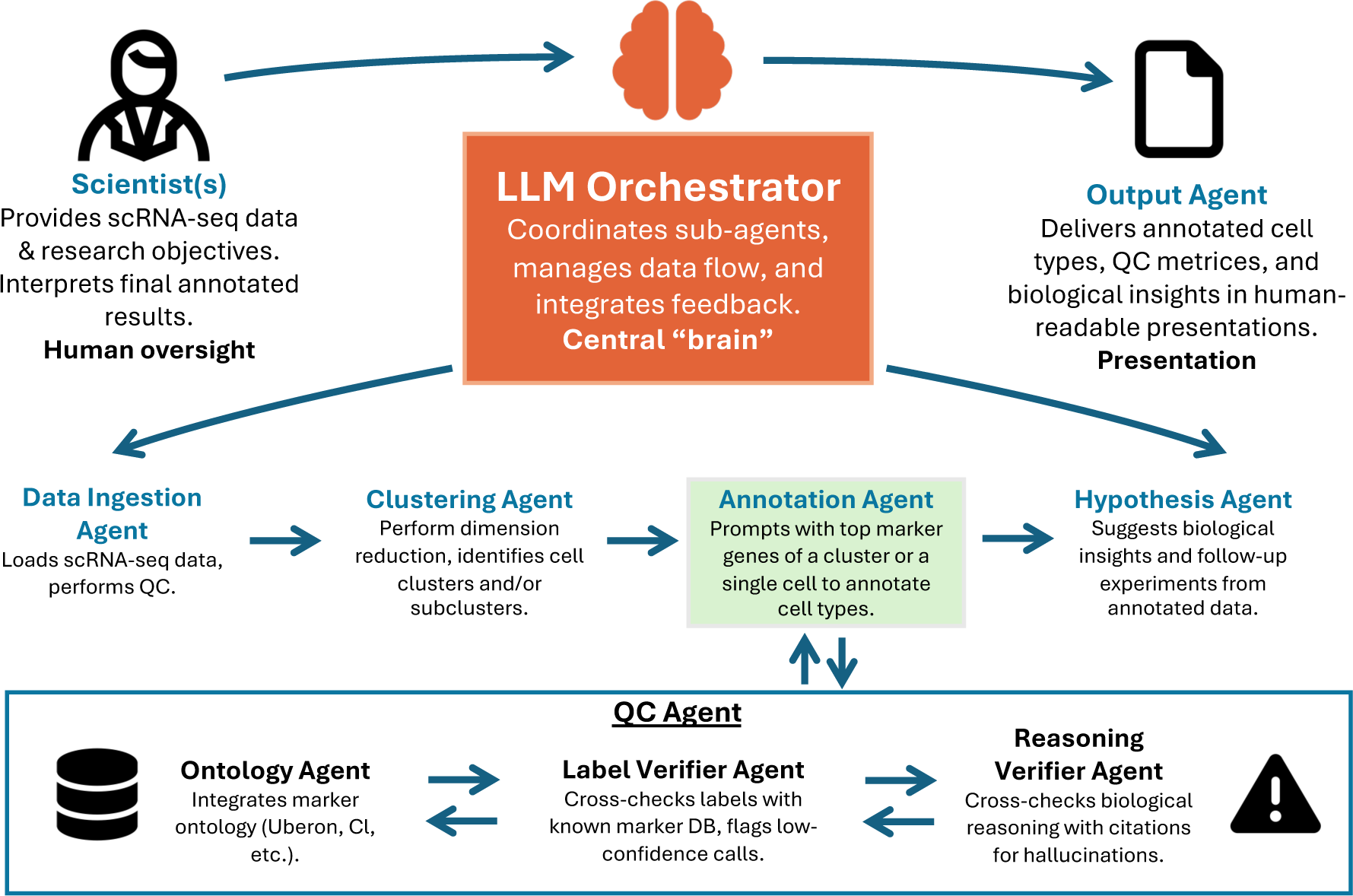
Proposed multi-agent reasoning LLM workflow for scRNAseq annotation. The LLM “brain” Orchestrator coordinates specialized sub-agents, each handling a distinct step of the scRNAseq bioinformatics pipeline from data ingestion and clustering to annotation and quality control. The Data Ingestion Agent loads raw scRNAseq data and performs initial checks; the Clustering Agent partitions cells into groups; and the Annotation Agent uses marker-gene prompts to assign preliminary labels. A QC layer, supported by the Ontology Agent and various verifier agents, cross-references known marker databases to flag potential hallucinations. Finally, the Output Agent consolidates refined annotations, metrics, and summary reports for researchers’ review. This orchestrated setup ensures transparency, modularity, and flexibility in automating the cell type annotation process.

## 4. Methods

### 4.1. Benchmarking datasets

#### 4.1.1. Cluster-level data

For the cluster-level analysis, we utilized the data curated by *Hou et al. (2024)*^7^, a study which assessed GPT-4’s ability to annotate single-cell clusters based on marker gene information. The data had been derived from a comprehensive list of human and mouse scRNAseq datasets spanning a wide range of tissues (Fig. 2A). We downloaded the annotated data containing human and mouse clusters (excluding non-model mammals) through the GitHub repo associated with the paper: https://github.com/Winnie09/GPTCelltype_Paper/blob/master/anno/compiled/all.csv. In the study, each cluster was assigned a set of 10 marker genes identified by a Wilcoxon-based differential expression analysis. We used these top 10 marker genes per cluster to prompt the LLMs in a zero-shot manner. If metadata (e.g., tissue name and/or disease) was available, we appended it to the prompt, as described below.

#### 4.1.2. Single-cell level data

##### In-Domain scTab datasets

We obtained the dataset from the scTab publication^14^, encompassing both validation and test pools of curated single cells. Specifically, we downloaded the data through the training data checkpoint from https://github.com/theislab/scTab/tree/devel using this specific link: https://pklab.med.harvard.edu/felix/data/merlin_cxg_2023_05_15_sf-log1p.tar.gz. To limit computational and API overhead, we randomly subsampled 10,000 cells from both the validation and the test sets for our benchmarking. We call these two pools of 10,000 cells *in-domain*.

##### Out-Of-Domain (OOD) random dataset

To evaluate generalizability beyond the training distribution of scTab, we curated a subset of scRNA-seq data from CellxGene^44^ with the following characteristics: human, primary (the study that originally generated the dataset), non-diseased, and added to the database after May 15^th^ 2023 – the training cutoff date for both scTab and scGPT. We call such data *Out-Of-Domain (OOD)*. A custom Python script first built an index of new unique cell-specific identifiers absent from the 2023-05-15 release and present only in 2024-07-01 release, then downloaded those cells in chunked form, storing the data in partitioned .h5ad files. After assembling all downloaded cells, we filtered out the cells labeled as unknown cell type and randomly subsampled 10,000 cells, reflecting the random *OOD* pool of normal tissues from CellxGene that scTab’s training and testing had not yet encountered. All cells passed our QC assessment.

##### OOD balanced tissue dataset

Beyond random sampling, we also built a “balanced” *OOD* dataset of 10,000 cells that more evenly covered multiple tissue types. Specifically, from the same post–2023-05-15 normal data obtained as explained above, we chose eight representative datasets of interest, as assessed by three criteria: i) a relatively high number of unique cell types profiled; ii) a relatively low fraction of cells labeled as unknown; iii) a relatively well-studied tissue. We randomly drew 1,250 cells from each tissue dataset, totaling 10,000 cells (see Supplementary Table S2 for an overview of the datasets we chose from). The tissues we chose were blood, peripheral region of retina, breast, lung, trachea, kidney, pancreas and cerebellum. For the lung dataset, 382 of the 1,250 cells had no match of the ground truth label in Cell Ontology and were therefore excluded from evaluations.

### 4.2. Prompting

We crafted prompt templates that encode key metadata (*e.g.*, tissue of origin, disease status) and a ranked gene list (either cluster-level marker genes obtained via differential expression or single-cell–level top highly expressed genes). We tested both a long prompt, which encouraged a thorough rationale, as well as a shorter variant, which requested a briefer answer.

#### 4.2.1. Cluster-level

We used a Python script to create prompts taking as input a CSV file in which each row described one cluster, including the top 10 marker genes determined by the differential expression analysis performed by *Hou et al. (2024)*^7^, as well as relevant metadata (*e.g.* tissue, disease status, etc.). The script outputted a line-delimited JSON file suitable for ingestion by downstream LLM pipelines.

We constructed both a long prompt and a short prompt. Below is the exact text used for the long prompt:

“You are an expert in single-cell biology.\n\n”

“Below is metadata for one cluster of cells along with its marker genes:\n”

“Tissue: [TISSUE_PLACEHOLDER]\n”

“Disease: [DISEASE_PLACEHOLDER]\n”

“Development stage: [DEVELOPMENT_STAGE_PLACEHOLDER]\n”

“Marker genes: [GENE_LIST]\n\n”

“Please identify what cell type this might be, as granular and accurate as possible.\n”

“At the end of your response, strictly place the final lines in this format:\n\n”

“Cell type: X\n”

Conversely, the short prompt added the instruction to “keep your response concise and clear”, while otherwise preserving the exact same structure:

“Please identify what cell type this might be, as granular and accurate as possible.\n”

“Keep your response concise and clear.\n”

“At the end of your response, strictly place the final lines in this format:\n\n”

“Cell type: X\n”

#### 4.2.2. Single-cell level

Below is the skeleton of our long prompt for querying LLMs to annotate a single cell:

“You are an expert in single-cell biology.\n\n”

“Below is metadata for one cell, followed by a list of its genes in descending expression:\n”

“Tissue: [TISSUE]\n”

“Disease: [DISEASE]\n”

“Development stage: [DEVELOPMENT_STAGE]\n”

“Genes: [GENE_LIST]\n\n”

“Please identify what cell type this might be, as granular and accurate as possible.\n”

“At the end of your response, strictly place the final lines in this format:\n\n”

“Cell type: X\n”

As in the case of the cluster-level annotations, the short prompt contained the additional instruction: “Keep your response concise and clear.”

A third variant of prompt incorporated a user-provided list of cell type labels from which the LLM was tasked to choose, effectively enforcing the model to act like a multi-class classifier. In our experiments, we used two different sets of classifier labels: the labels curated by scTab, and the labels provided by scGPT. The relevant lines added to the prompt were:

“Here is a list of all possible cell types you must choose from:\n”

“[MULTILINE_STRING_OF_CELL_TYPES]\n\n”

“Please pick the single best matching cell type from this list.\n”

We used multiple values of N (the number of top expressed genes) for prompting: 5, 10, 25, 50, 100, 200, 500, 1000, 2000, as well as all the non-zero expressing genes. We settled on top 500 most highly expressed genes for final input prompts, as they achieved top accuracy and Macro-F1 score.

### 4.3. Performance evaluation

To evaluate the performance of the tested models, we considered as ground truth the labels curated and proposed by the original dataset. In order to compare model predictions with ground truth labels, we first harmonized both predicted and ground truth labels to official Cell Ontology (CL) terms, as done by *Hou et al. (2024)*^7^. This step was necessary because, unlike multi-class classifier models like scTab, where the predicted cell type labels are standardized to CL terms, LLM outputs could vary by case, wording, or description, even if conceptually in agreement with the ground truth label. After the harmonization process, we removed the cells with no match (empty match) of the ground truth label with the OLS database, specifically 94 cells from the *in-domain* validation pool, 87 cells from the *in-domain* test pool, 31 cells from the *OOD* random dataset, and 382 lung cells from the *OOD* balanced tissue dataset. Next, given the unambiguous ground truth label and unambiguous predicted label, we applied the scTab^14^ framework as described in their GitHub page (https://github.com/theislab/scTab/tree/devel). Specifically, a match was achieved (return TRUE) if the predicted label was either identical (exact match) or a Cell Ontology descendant (predicted subtype) of the ground truth label. Conversely, if the ground truth label was a child of the predicted label or the classes shared no direct parent–child relationship, the result was incorrect (return FALSE). In other words, the predicted labels were not penalized for higher granularity than ground truth labels. Finally, we compiled the cell-level binary TRUE/FALSE summary statistics to compute overall prediction accuracies and Macro-F1 scores for each dataset and for each model, as detailed below.

#### 4.3.1. Accuracy

We computed the accuracy score as follows:

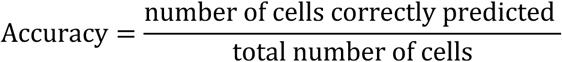

#### 4.3.2 Macro-F1

To obtain the macro-averaged F1 score, for every ground-truth cell-type label ℓ, we counted:

𝑻𝑷_𝓵_: cells whose ground-truth label is ℓ and were correctly predicted as either ℓ or an ontology descendant of ℓ (label TRUE),

𝑭𝑵_𝓵_: cells whose ground-truth label is ℓ and were incorrectly predicted (label FALSE),

𝑭𝑷_𝓵_: cells whose ground-truth label is not ℓ but whose predicted label is ℓ (label FALSE).

We compute the per-label precision and recall metrics:

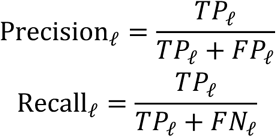

And the per-label F1 score:

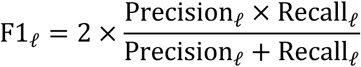

The Macro-F1 score is the average over all L ground-truth labels:

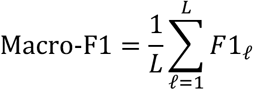

#### 4.3.3. Weighted-F1

Weighted-F1 score gives each cell type label a weight proportional to its relative frequency in the dataset. Therefore, if for every ground truth label ℓ we define 𝑛_ℓ_= 𝑇𝑃_ℓ_ + 𝐹𝑁_ℓ_ as the number of cells with the true label ℓ, then

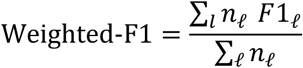

#### 4.3.4. Bootstrapped Confidence Intervals

To complement the point estimates for the single-cell *in-domain* and random *OOD* evaluations, we report 95 % confidence intervals (CI) around accuracy and macro-F1 values, obtained through a percentile bootstrap procedure. For each dataset consisting of *n* cells (*in-domain* validation 9,906 cells, *in-domain* test 9,913 and *OOD* random 9,969), we drew 10,000 bootstrap resamples, each consisting of *n* cell indices selected with replacement from the original index set. The same sampled indices were applied to the ground-truth vector and to the prediction vectors, to generate the bootstrapped evaluation vectors for all models. On every bootstrapped evaluation vector, we recomputed the metric of interest (accuracy or Macro-F1) for each model exactly as described above. For each model, we took the 2.5-th and 97.5-th percentiles of the 10,000 bootstrap replicates as the lower and upper bounds of the 95 % CI. As the *OOD* balanced tissue dataset was composed specifically of eight tissues with only 1,250 cells/tissue, we didn’t bootstrap CIs for this dataset.

#### 4.3.5. Pair-wise statistical significance testing

To determine whether performance differences between models were significantly different than expected by sampling variability, we carried out pair-wise significance tests separately for each dataset and for each metric (accuracy and Macro-F1). All p-values were adjusted for multiple testing by the Benjamini– Hochberg^52^ false-discovery-rate (FDR) procedure with α = 0.05.

##### Accuracy

For every pair of models, we formed a 2 × 2 contingency table:

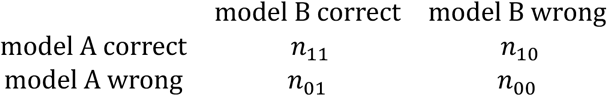

where 𝑛_10_ denotes cells called as correct by model A but wrong by model B and *vice versa* for 𝑛_01_. Under the null hypothesis that the two models have identical accuracy, the number of discordant pairs favoring model A, 𝑛_10_, follows a *Binomial (N, 0.5)* distribution with *N* = 𝑛_01_ + 𝑛_10_. We therefore applied the exact McNemar^53^ test (two-sided, binomial formulation) implemented in statsmodels^54^ 0.14.4. Given a total of 𝑘 models tested, the resulting exact p-values were FDR-adjusted across the 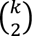 model pairs within the same dataset.

##### Macro-F1

Because macro-F1 is not a per-cell statistic, we assessed significance via a paired non-parametric bootstrap^55^ with 10,000 resamples. For each resample, we first sampled *n* cell indices with replacement from the dataset (where *n* is the number of evaluation cells) as described at the beginning of this subsection. We then recomputed the Macro-F1 score for every model on the resample and recorded the difference for each pair of models A and 𝐵 as Δ = Macro − F1_*_ − Macro − F1_+_. The empirical distribution of Δ over the 10, 000 replicates approximates its sampling distribution. A two-sided p-value was obtained as:

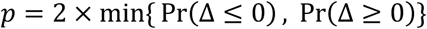

Bootstrap p-values for all model pairs were then FDR-corrected as described above.

### 4.4. scGPT

We employed the scGPT^15^ single-cell foundation model for benchmarking cell type annotation at the single-cell level. Our approach utilized the scGPT package and followed established procedures from the scGPT GitHub tutorial for zero-shot annotation via embedding similarity search, available here: https://github.com/bowang-lab/scGPT/blob/main/tutorials/Tutorial_Reference_Mapping.ipynb. The input data was provided as AnnData objects (.h5ad files) containing raw gene expression counts. Initial preprocessing involved filtering the genes present in the input data to retain only those included in the scGPT model’s vocabulary, ensuring compatibility between the input data features and the pre-trained scGPT model. Subsequently, we identified the top 3,000 highly variable genes (HVGs) within the filtered gene set using the scanpy.pp.highly_variable_genes function with the flavor=’seurat_v3’ setting from the scanpy^56^ Python package. The AnnData object was then subsetted to include only these 3,000 HVGs for downstream processing.

Cell embeddings were generated using the scgpt.tasks.embed_data function from the scGPT library, applying the pre-trained scGPT whole-human model (checkpoint-0623 from scGPT GitHub) to obtain a low-dimensional representation for each cell.

Zero-shot cell type annotation was performed using a nearest-neighbor approach based on these cell embeddings, which is the method recommended in scGPT tutorials. We utilized a pre-computed FAISS index built from reference cell embeddings derived from the CellXGene atlas (downloaded following scGPT GitHub instructions). For each cell in our input dataset, we queried the FAISS index to find the 50 nearest neighbors (k=50) within the reference atlas embedding space. The cell type labels associated with these 50 neighbors were retrieved from the reference metadata. A final cell type prediction for each input cell was determined by majority voting among the labels of its 50 nearest neighbors, using the voting function adapted from the scGPT repository’s utility scripts. The resulting single-cell level predictions were saved alongside ground truth labels for subsequent evaluation.

### 4.5. scTab

We also employed the scTab^14^ multi-class classification model for benchmarking cell type annotation at the single-cell level, following the scTab GitHub tutorials available here: https://github.com/theislab/scTab. Input data consisted of AnnData objects (.h5ad files) with raw gene expression counts in the .X attribute. Consistent with the scTab protocol, the feature space was first streamlined. Genes present in the input data were filtered and ordered to match the reference gene set used for model training. This step ensured that the input matrix dimensions and gene order align precisely with the model’s expectations. The streamline_count_matrix function from the cellnet library was utilized for this alignment^14^. Following gene alignment, the count matrix underwent normalization. Each cell’s expression profile was scaled to a total count of 10,000, followed by a log(x+1) transformation. This normalization method is standard practice for the scTab model.

For inference, we used the same pre-trained scTab checkpoint from their official GitHub tutorial (scTab-checkpoints/scTab/run5/val_f1_macro_epoch=41_val_f1_macro=0.847.ckpt). Corresponding model hyperparameters were loaded from the associated hparams.yaml file within the same directory. The TabNet classifier architecture was initialized using these parameters and loaded with the extracted state dictionary from the checkpoint. Inference was performed on the normalized data using PyTorch DataLoader for batch processing, leveraging GPU acceleration. The model outputs raw logits, from which the predicted class index (representing the cell type) was determined using an argmax operation. Finally, these numeric prediction indices were mapped to human-readable cell type labels using the official mapping file (merlin_cxg_2023_05_15_sf-log1p_minimal/categorical_lookup/cell_type.parquet). The resulting single-cell level predictions were saved alongside ground truth labels for subsequent evaluation.

### 4.6. Cell2Sentence-Scale-1B

We followed the exact instructions on the cell2sentence official GitHub page tutorials for zero-shot cell type annotations, especially tutorials 0 and 6 at the following link as of June 10^th^ 2025 : https://github.com/vandijklab/cell2sentence/tree/master/tutorials. Explicitly, we did not perform the simple QC steps recommended in the tutorial 0, as the data we had were already QCed. In addition, we did not have “batch_condition” metadata for our samples, and we used the “sex” metadata information when available. We used the newest release of cell2sentence-scale model C2S-Scale-Pythia-1b-pt (https://huggingface.co/vandijklab/C2S-Scale-Pythia-1b-pt) in our evaluation, as this was the only new available model released by the cell2sentence-scale team at the time of testing. We used the default number of 200 genes in the prompt, as shown in the GitHub tutorials.

## 5. Data and code availability

All processed data, prompts, responses and scripts are available at: https://github.com/cristea-lab/BioReasoning_Paper.

## 6. Financial statement and API use

At the time of this project, API calls for DeepSeek-R1-0528 from the DeepSeek Platform were $0.14/1M tokens input (cache hit), $0.55/1M tokens input (cache miss), and $2.19/1M tokens output. For DeepSeek-V3 released on 12/26/2024, as well as DeepSeek-V3-0324, API prices from the DeepSeek Platform were $0.07/1M tokens input (cache hit), $0.27/1M tokens output (cache miss), and $1.10/1M tokens output. For GPT-4o, we used the API service provided by OpenAI, which was priced at $2.50/1M tokens input, $1.25/1M tokens cached input, and $10/1M tokens output. For DeepSeek-R1-0528, this study used 832 million tokens (353 million input tokens, 203 million reasoning tokens, and 276 million completion tokens) across 144,520 API calls.

## 7. Author contributions

X.W. and S.C. conceived the study. R.T. and X.W. conducted the analyses. X.W., R.T., B.W., and S.C interpreted the results. S.C. supervised the study. All authors wrote and approved the final manuscript.

