## Supplementary Figures for "Biological Reasoning with Reinforcement Learning through Natural Language Enables Generalizable Zero-Shot Cell Type Annotations"

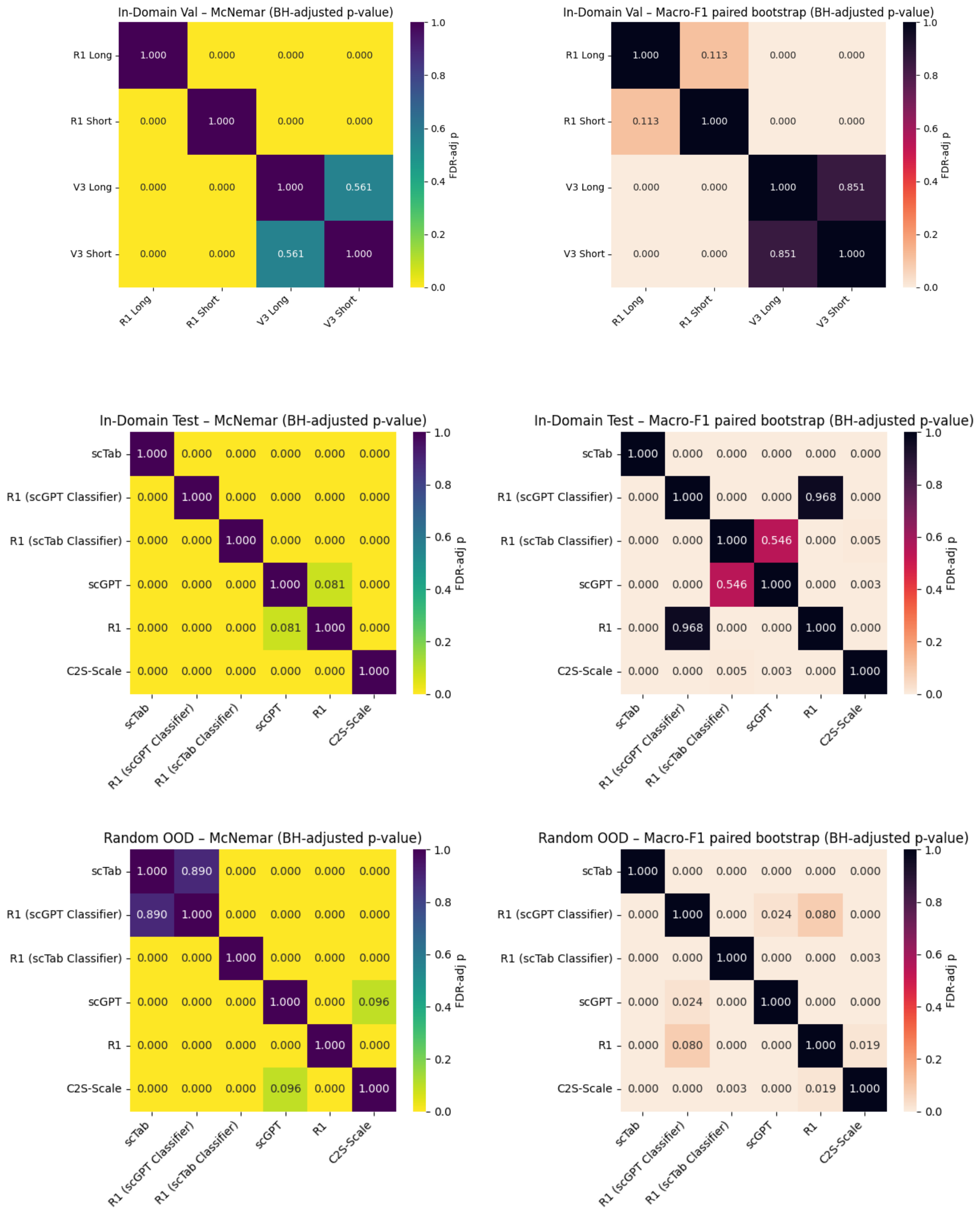
In-Domain Val – McNemar (BH-adjusted p-value)
R1 Long: 1.000 0.000 0.000 0.000
R1 Short: 0.000 1.000 0.000 0.000
V3 Long: 0.000 0.000 1.000 0.561
V3 Short: 0.000 0.000 0.561 1.000
R1 Long R1 Short V3 Long V3 Short
FDR-adj p
In-Domain Val – Macro-F1 paired bootstrap (BH-adjusted p-value)
R1 Long: 1.000 0.113 0.000 0.000
R1 Short: 0.113 1.000 0.000 0.000
V3 Long: 0.000 0.000 1.000 0.851
V3 Short: 0.000 0.000 0.851 1.000
R1 Long R1 Short V3 Long V3 Short
FDR-adj p
In-Domain Test – McNemar (BH-adjusted p-value)
scTab: 1.000 0.000 0.000 0.000 0.000 0.000
R1 (scGPT Classifier): 0.000 1.000 0.000 0.000 0.000 0.000
R1 (scTab Classifier): 0.000 0.000 1.000 0.000 0.000 0.000
scGPT: 0.000 0.000 0.000 1.000 0.081 0.000
R1: 0.000 0.000 0.000 0.081 1.000 0.000
C2S-Scale: 0.000 0.000 0.000 0.000 0.000 1.000
scTab R1 (scGPT Classifier) R1 (scTab Classifier) scGPT R1 C2S-Scale
FDR-adj p
In-Domain Test – Macro-F1 paired bootstrap (BH-adjusted p-value)
scTab: 1.000 0.000 0.000 0.000 0.000 0.000
R1 (scGPT Classifier): 0.000 1.000 0.000 0.000 0.968 0.000
R1 (scTab Classifier): 0.000 0.000 1.000 0.546 0.000 0.005
scGPT: 0.000 0.000 0.546 1.000 0.000 0.003
R1: 0.000 0.968 0.000 0.000 1.000 0.000
C2S-Scale: 0.000 0.000 0.005 0.003 0.000 1.000
scTab R1 (scGPT Classifier) R1 (scTab Classifier) scGPT R1 C2S-Scale
FDR-adj p
Random OOD – McNemar (BH-adjusted p-value)
scTab: 1.000 0.890 0.000 0.000 0.000 0.000
R1 (scGPT Classifier): 0.890 1.000 0.000 0.000 0.000 0.000
R1 (scTab Classifier): 0.000 0.000 1.000 0.000 0.000 0.000
scGPT: 0.000 0.000 0.000 1.000 0.000 0.096
R1: 0.000 0.000 0.000 0.000 1.000 0.000
C2S-Scale: 0.000 0.000 0.000 0.096 0.000 1.000
scTab R1 (scGPT Classifier) R1 (scTab Classifier) scGPT R1 C2S-Scale
FDR-adj p
Random OOD – Macro-F1 paired bootstrap (BH-adjusted p-value)
scTab: 1.000 0.000 0.000 0.000 0.000 0.000
R1 (scGPT Classifier): 0.000 1.000 0.000 0.024 0.080 0.000
R1 (scTab Classifier): 0.000 0.000 1.000 0.000 0.000 0.003
scGPT: 0.000 0.024 0.000 1.000 0.000 0.000
R1: 0.000 0.080 0.000 0.000 1.000 0.019
C2S-Scale: 0.000 0.000 0.003 0.000 0.019 1.000
scTab R1 (scGPT Classifier) R1 (scTab Classifier) scGPT R1 C2S-Scale
FDR-adj p

**Supplementary Figure S1:**

Heatmaps of statistical significance tests for the *in-domain* validation (top), *in-domain* test (middle), and *OOD* random (bottom) datasets for accuracy and Macro-F1 score, as described in Methods. For each dataset, we report two heatmaps: 1) Left-hand panels (yellow-green colormap): Benjamini-Hochberg-adjusted[1] p-values from the exact McNemar[2] test applied to per-cell accuracy. 2) Right-hand panels (orange–purple colormap): Benjamini-Hochberg-adjusted p-values from a paired, non-parametric bootstrap[3] test on Macro-F1 score with 10,000 bootstrap resamples.

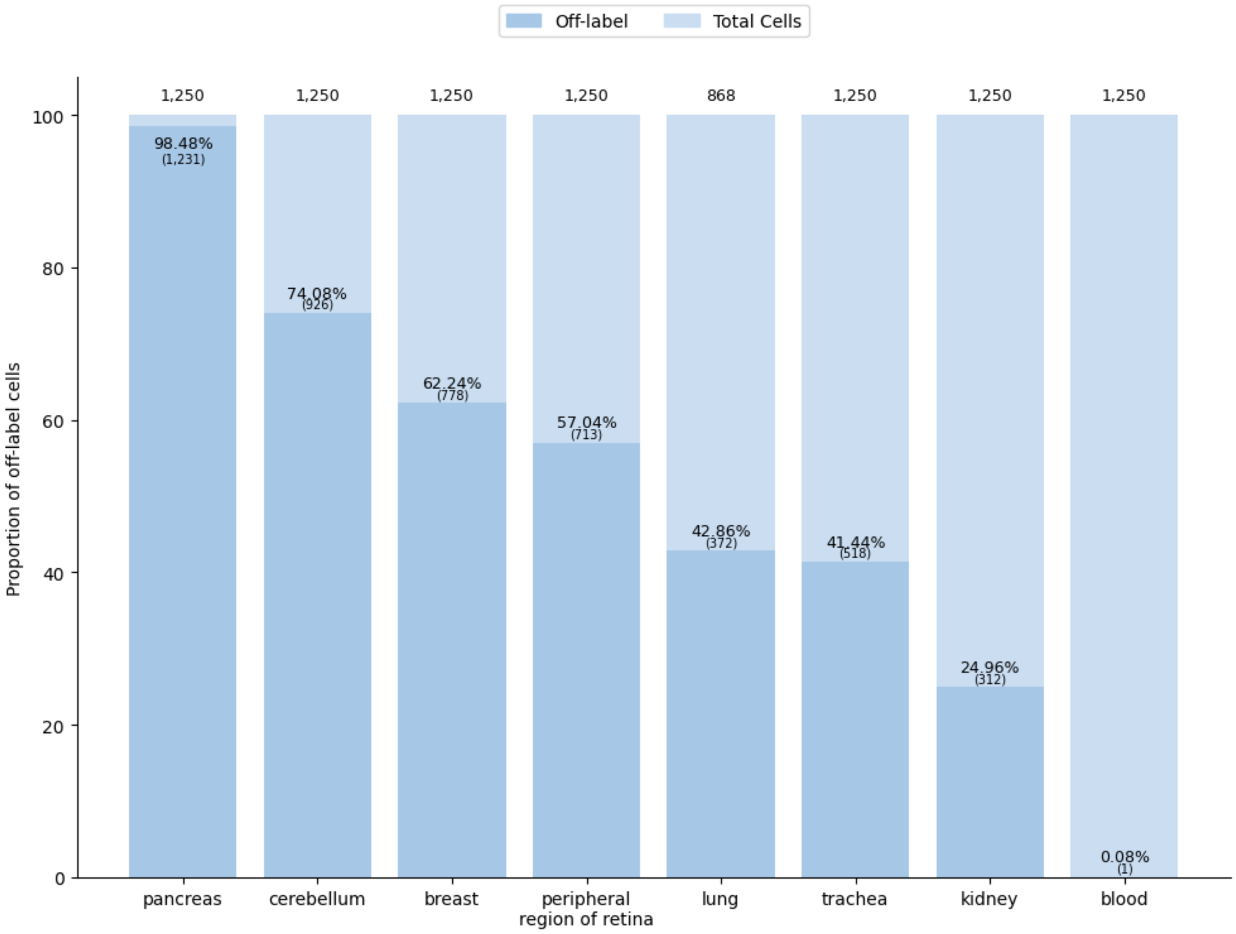

Off-label
Total Cells
Proportion of off-label cells
1,250
1,250
1,250
1,250
868
1,250
1,250
1,250
98.48%
(1,231)
74.08%
(926)
62.24%
(778)
57.04%
(713)
42.86%
(372)
41.44%
(518)
24.96%
(312)
0.08%
(1)
0
20
40
60
80
100
pancreas
cerebellum
breast
peripheral region of retina
lung
trachea
kidney
blood

**Supplementary Figure S2:**
Bar plot showing the proportion of off-scTab-label[4] cells by tissue types in the *OOD* balanced tissue dataset, with shades of color showing whether they were outside the labels employed by scTab (darker color indicates off-label). Percentages show the proportion of off-label cells for each tissue type, and absolute numbers show the number of cells in each category accordingly.

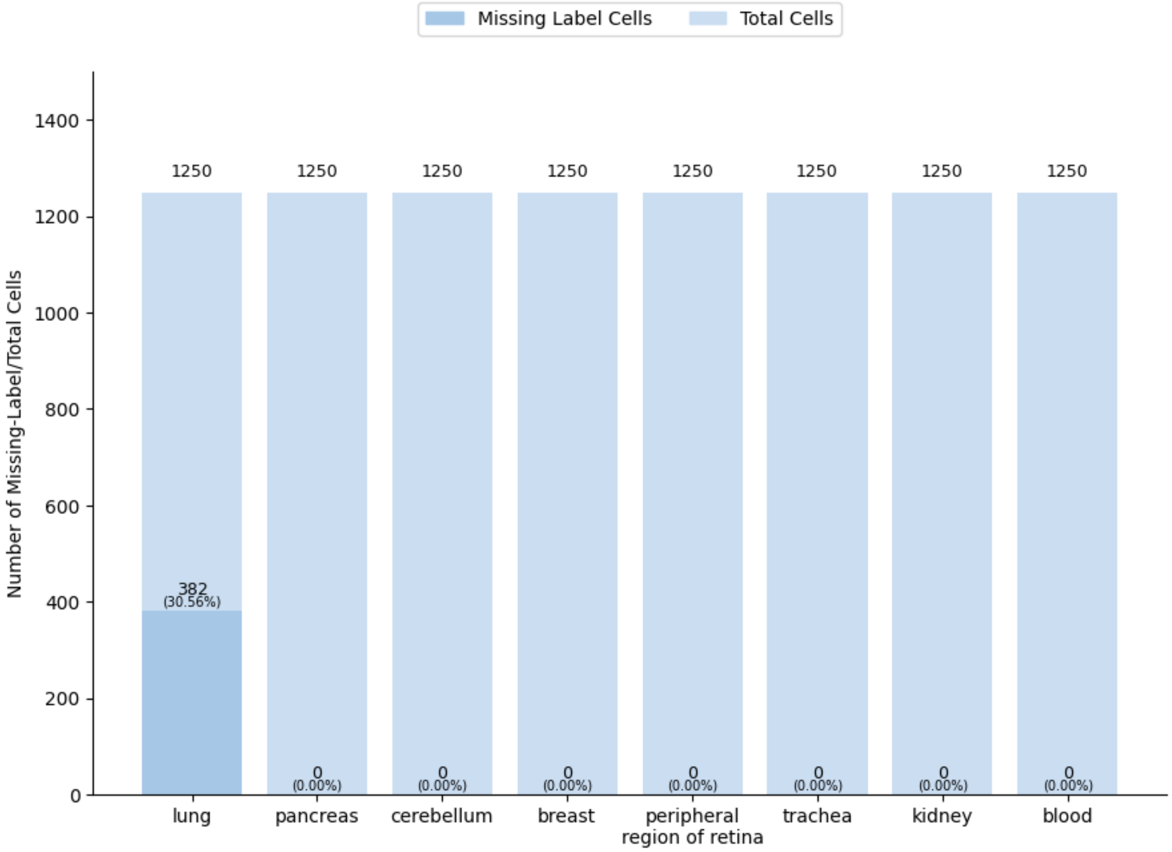
Missing Label Cells
Total Cells
Number of Missing-Label/Total Cells
1400
1200
1000
800
600
400
200
0
1250
1250
1250
1250
1250
1250
1250
1250
382
(30.56%)
0
(0.00%)
lung
pancreas
cerebellum
breast
peripheral region of retina
trachea
kidney
blood

**Supplementary Figure S3:**
Bar plot showing the proportion of missing-label cells by tissue types in the *OOD* balanced tissue dataset, with shades of color showing whether they don't have a ground truth cell type from the Cell Ontology Database[5] (darker color indicates missing-label). Percentages show the proportion of missing-label cells for each tissue type, and absolute numbers show the number of cells in each category accordingly.
